# Redefining the *PTEN* Promoter: Identification of Two Upstream Transcription Start Regions

**DOI:** 10.1101/2021.04.23.441162

**Authors:** Dennis Grencewicz, Todd Romigh, Stetson Thacker, Ata Abbas, Ritika Jaini, Donal Luse, Charis Eng

**Affiliations:** Genomic Medicine Institute, Lerner Research Institute, Cleveland Clinic, Cleveland, OH, 44195, USA; Department of Cardiovascular and Metabolic Sciences, Lerner Research Institute, Cleveland Clinic, Cleveland, OH 44195, USA; Department of Molecular Medicine, Cleveland Clinic Lerner College of Medicine, Case Western Reserve University, Cleveland, Ohio 44106, U.S.A.; Division of Hematology and Oncology, Department of Medicine, Case Western Reserve University, Cleveland, OH 44106; Developmental Therapeutics Program, CASE Comprehensive Cancer Center, Case Western Reserve University School of Medicine, Cleveland, OH 44106, USA; Germline High Risk Focus Group, CASE Comprehensive Cancer Center, Case Western Reserve University School of Medicine, Cleveland, OH 44106, U.S.A.; Center for Personalized Genetic Healthcare, Cleveland Clinic Community Care and Population Health, Cleveland, Ohio 44195, U.S.A.; Department of Genetics and Genome Sciences, Case Western Reserve University School of Medicine, Cleveland, Ohio 44106, U.S.A.

## Abstract

Germline mutation of *PTEN* is causally observed in Cowden syndrome (CS) and is one of the most common genetic causes of autism spectrum disorder (ASD). However, the majority of individuals who present with CS-like clinical features are found to be *PTEN-*mutation negative. Reassessment of *PTEN* promoter regulation may help explain abnormal *PTEN* dosage, as only the minimal promoter and coding regions are currently included in diagnostic *PTEN* mutation analysis. We reanalyzed the architecture of the *PTEN* promoter using next-generation sequencing datasets. Specifically, run-on sequencing assays identified two additional TSRs at −2052 and −1907 basepairs from the start of PTEN, thus redefining the *PTEN* promoter and extending the *PTEN* 5’UTR. The upstream TSRs described are active in cancer cell lines and human cancer and normal tissue. Further, these TSRs can produce novel *PTEN* transcripts due to the introduction of new splice sites. Evaluation of transcription factor binding specific to the upstream TSRs shows overrepresentation of TFs involved in inflammatory processes. Together, these data suggest that potentially clinically relevant promoter variants upstream of the known promoter may be overlooked in indivduals considered *PTEN* germline mutation-negative and may also explain lack of PTEN expression in sporadic neoplasias without *PTEN* somatic structural defects.

## INTRODUCTION

The tumor suppressor gene *PTEN* encodes a dual-specificity phosphatase and tensin homolog (1–3) implicated, via germline mutation, in overgrowth and neurodevelopmental phenotypes, such as cancer and autism spectrum disorder (ASD; 4, 5) and, via somatic mutation, in numerous sporadic malignancies (6–10). In its canonical role, PTEN antagonizes the PI3K/AKT/mTOR signaling pathway through dephosphorylation of phosphatidylinositol (3,4,5)-triphosphate (PIP_3_) to phosphatidylinositol 4,5-bisphosphate (PIP_2_; 11). Because of *PTEN*’s additional role in cell cycle control and cell proliferation, perturbations in *PTEN* expression can contribute to tumorigenesis (4). The molecular diagnosis *PTEN* Hamartoma Tumor Syndrome (PHTS) unifies subsets of any clinical presentation caused by germline *PTEN* mutations. These syndromes include Cowden Syndrome (CS), Bannayan-Riley-Ruvalcaba Syndrome (BRRS), and Proteus Syndrome (PS; (12–14). PHTS individuals have an increased lifetime risk of developing primary and secondary cancers (4). *PTEN* mutations are also strongly associated with neurodevelopmental disorders (NDDs), including autism spectrum disorder (ASD); between 7-27% of individuals with macrocephalic ASD will carry a germline *PTEN* mutation (5, 15–17), making *PTEN* one of the most common ASD-predisposition genes.

Interestingly, 15-75% percent of CS/CS-like and 40% of BRRS patients are clinically negative for germline *PTEN* mutations (12). This missing genetic etiology is an area of active research. Because decreases in the tumor suppressor gene (TSG) gene dosage may drive phenotypic change (18–20), the *PTEN* promoter, as currently defined, is also included in routine clinical sequencing for individuals with suspected PHTS. Prior work has demonstrated that even small decreases in *Pten* expression associate with increased breast tumorigenesis, establishing a precedent for continued assessment of *PTEN* promoter dysregulation and other mechanisms that regulate *PTEN* expression (21). Additionally, *PTEN* promoter variants may have functional effects beyond suppressing expression. Many constitutively expressed TSGs are subject to multiple regulatory mechanisms that can operate tissue-specifically (22), including utilization of different transcription start regions (TSRs; 22, 23). Reports of multiple PTEN proteoforms and alternatively spliced isoforms also suggest that *PTEN* promoter variants may be relevant to PHTS phenotypes through these additional mechanisms (24–31). Notably, particular PTEN proteoforms are enriched in breast and renal-cell carcinoma (24, 27). However, much remains to be determined about non-canonical transcriptional and translational regulatory processes governing *PTEN* expression. Here, we hypothesize that a comprehensive reassessment of *PTEN’s* upstream region may reveal novel landmarks or regulatory achitecture that have heretofore been overlooked.

Multiple studies have attempted to uncover the missing genetic origins of disease in PHTS-like *PTEN* mutation-negative individuals by studying alternate *PTEN* promoter variants, regulatory domains, and transcription start sites (TSSs) in relation to CS and breast cancer (32–35). Currently, *PTEN* has one identified human promoter. The minimal *PTEN* promoter is defined by −958 to −821 of the translational start codon, containing approximately 10 putative TSSs, was defined by Sheng et al. (36), with the putative strongest transcription start site (MaxTSS) between −951 and −925 (37). A second *Pten* TSR has been described as a constitutive promoter in mice, which was mapped between −551 and −220. However, it remains questionable whether this region also acts as a TSR in humans (38). Additionally, there is an enhancer-box (E-Box) domain 1.2kb upstream of the Sheng promoter has been shown to bind USF1 and USF2 and can mediate *PTEN* transcriptional activation (34, 39). Currently, the accepted definition of the *PTEN* promoter spans form −1344 to −745 (32), indicating that any mutations upstream or downstream of the loosely defined Sheng promoter, such as near the E-box domain or within the region identified by Han and colleagues, are not assessed. This constrained promoter definition (Figure 1A) represents a potential gap in coverage for identifying variants that may have clinically relevant effects on *PTEN* transcription and subsequent phenotypes.

**Figure 1:**
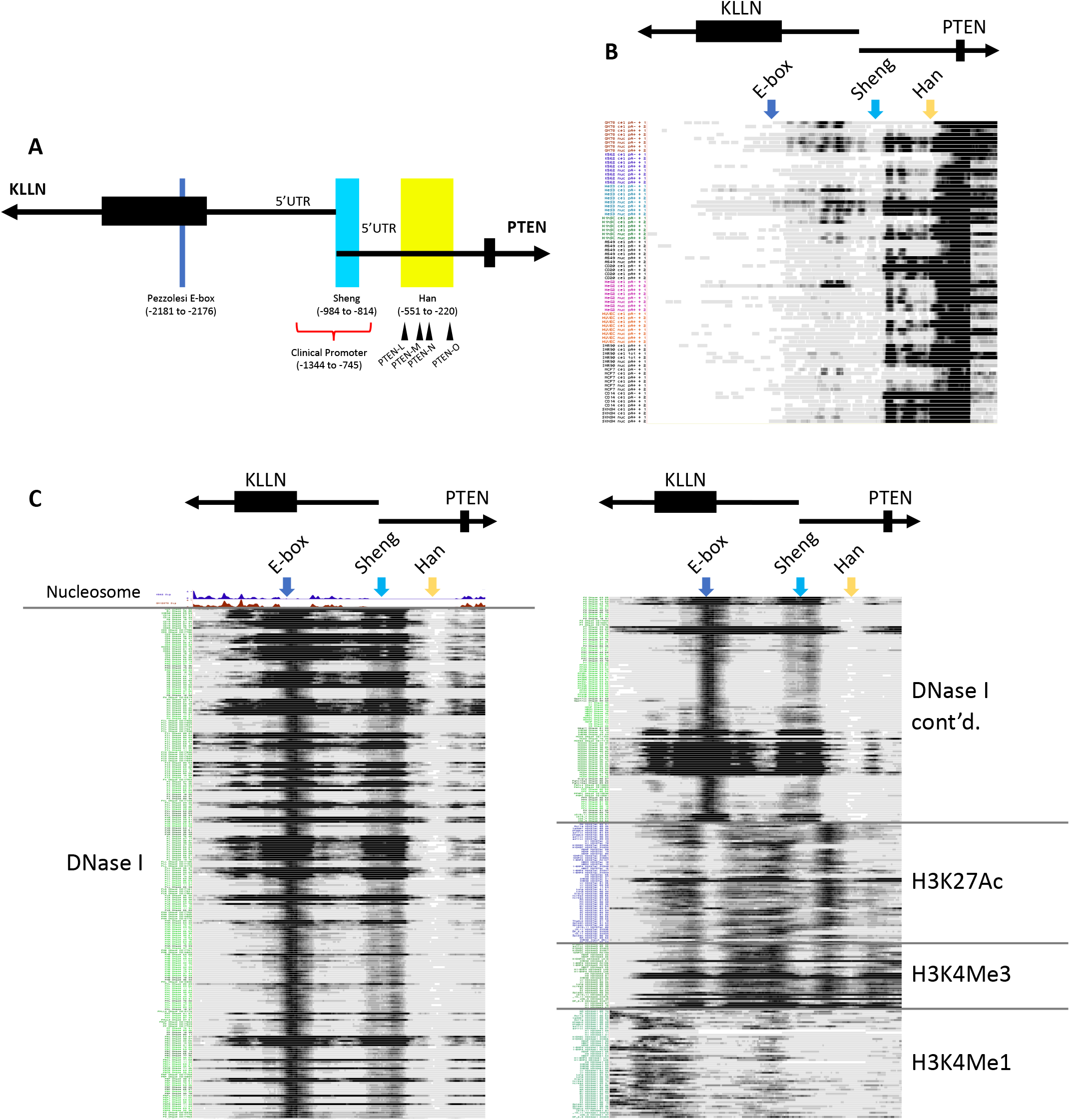
Genome browser data allude to transcription of *PTEN* upstream of its canonical promoter. (A) Schematic of known transcription regulatory regions of *PTEN* and locations of proteoform translation start sites. Nucleotide coordinates defined by distance upstream of the canonical AUG translation start site. (B) ENCODE 3 RNA-sequencing tracks for Tier 1 and 2 cell lines showing full cell and nucleus-specific analyses aligned to upstream of *PTEN* locus. Includes poly(a) +/-tracks per cell type, with replicates shown as “1” and “2”. (C) ENCODE 3 NGS data visualized through UCSC Genome Browser suggest multiple regions in *PTEN* upstream region may be involved with transcriptional regulation. Nucleosome tracks show nucleosome depletion throughout canonical promoter and E-box domain in K562 and GM12878 cell lines. DNase I Hypersensitivity tracks throughout all ENCODE 3 accessible tissue samples show open chromatin conformation Sheng promoter and near E-box domain. Promoter-associated histone marks (H3K27Ac and H3K4Me3) in human tissue samples from ENCODE show flanking pattern to E-box region and Sheng promoter. Enhancer-associated histone mark (H3K4Me1) generally depleted throughout upstream of *PTEN* locus.

Although previous studies provide foundational insights into *PTEN* transcriptional regulation, all were completed before the advent of next-generation sequencing (NGS) technologies. Many high-throughput assays have been developed that expand our understanding of global transcriptional regulation. Here, we present comprehensive alignments of multiple forms of NGS data (23, 40–45) using the UCSC Genome Browser (46), followed by empirical validation studies that led to identification of two novel TSRs for *PTEN*, upstream of the current clinically-relevant promoter region.

## MATERIALS AND METHODS

### Biological Data Resources

All breast and thyroid tissues were tumor/normal pairs from the Cooperative Human Tissue Network (CHTN). Cell lines included one patient derived lymphoblast cell line, four breast cell lines (MCF10A, ATCC CRL-10317; BT474, ATCC CRL-3247; MCF7, ATCC HTB-22, and MDA-MB-231, ATCC HTB-26), and four thyroid cell lines (Nthy-ori 3-1, ECACC 90011609; B-CPAP, ACC 273; FTC-236 ECACC 94060901; R082-W-1, ECACC 92030502), all acquired from ATCC. Lines were cultured with cell-specific media at 37°C and 5% CO_2_ using standardized protocol.

### Molecular Analyses

#### Total RNA Extraction

20-50 mg of tissue or cells were homogenized in 1 ml of Trizol (ThermoFisher Cat # 15596026) according to the manufacturer’s protocol in a TissueRuptor (Qiagen Cat # 9002755) homogenizer. The mixture was added to an RNeasy Column (Qiagen Cat # 74101) and processed according to manufacturer’s protocol including the on-column DNase I treatment (Qiagen Cat # 79254).

### Quantitative Reverse Transcription-PCR

3µg of RNA was reverse transcribed using SuperScript III kit and oligo(dT) primers (ThermoFisher Cat # 18080051) according the manufacturer’s protocol 100ng cDNA was amplified with AmpliTaq Gold™ 360 (Cat # 4398876), 5% GC Enhancer, and 0.2uM of each primer in a 25ul reaction. Forward primers were located within the *KLLN* coding sequence (CDS), *KLLN* 5’ untranslated region (UTR), or *PTEN* 5’UTR and paired with reverse primers located either in *PTEN* exon 5 or downstream of the *PTEN* canonical stop codon in exon 9. PCR Bands were resolved by running 18ul of the PCR product on a 1% agarose gel with ethidium bromide.

### Primers Utilized

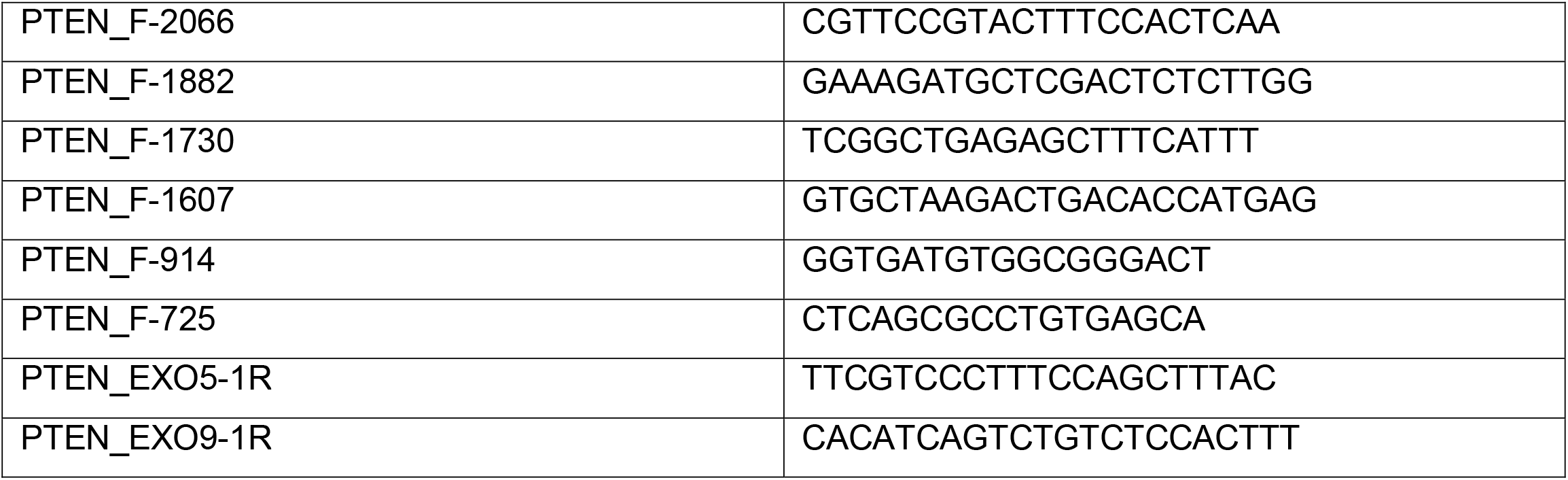

### 5’ Rapid Amplification of cDNA Ends

5’ ends of PTEN-specific transcripts were assessed in one patient derived lymphoblast cell line using the Takara SMARTer RACE kit (Cat # 634858) with a R-Ex4 PTEN 5’RACE gene-specific primer using the manufacturer designated protocol. PTEN-specific GSP used the following sequence: GATTACGCCAAGCTTGGCGGTGTCATAATGTCTTTCAGCAC.

### Cloning of PCR Products, Whole Cell PCR and Sequencing

Unpurified PCR products were directly cloned using StrataClone PCR Cloning Kit (Cat # 240205) according to manufacturer’s protocol. Colonies were picked and the cloned sequence amplified using AmpliTaq Gold™ 360 as described above but with standard T3 and T7 primers. Subsequent amplicons were Exo-SAP treated (Exo I, NEB Cat # M0293S and SAP, Affymetrix Cat # 78390) and Sanger sequenced.

### Computational Data Resources

#### Genome Browser Alignments

The UCSC Genome Browser was used to align custom and publicly available genome browser tracks (http://genome.ucsc.edu). Majority of tracks represent data collected for the ENCODE project (40, 41, 43– 45). Custom tracks generated using BLAT for 5’RACE-seq experiments (Figure 3A). Nanopore sequencing was visualized using IGV viewer (Supplemental Figure 1).

#### Links and Sources for UCSC Tracks

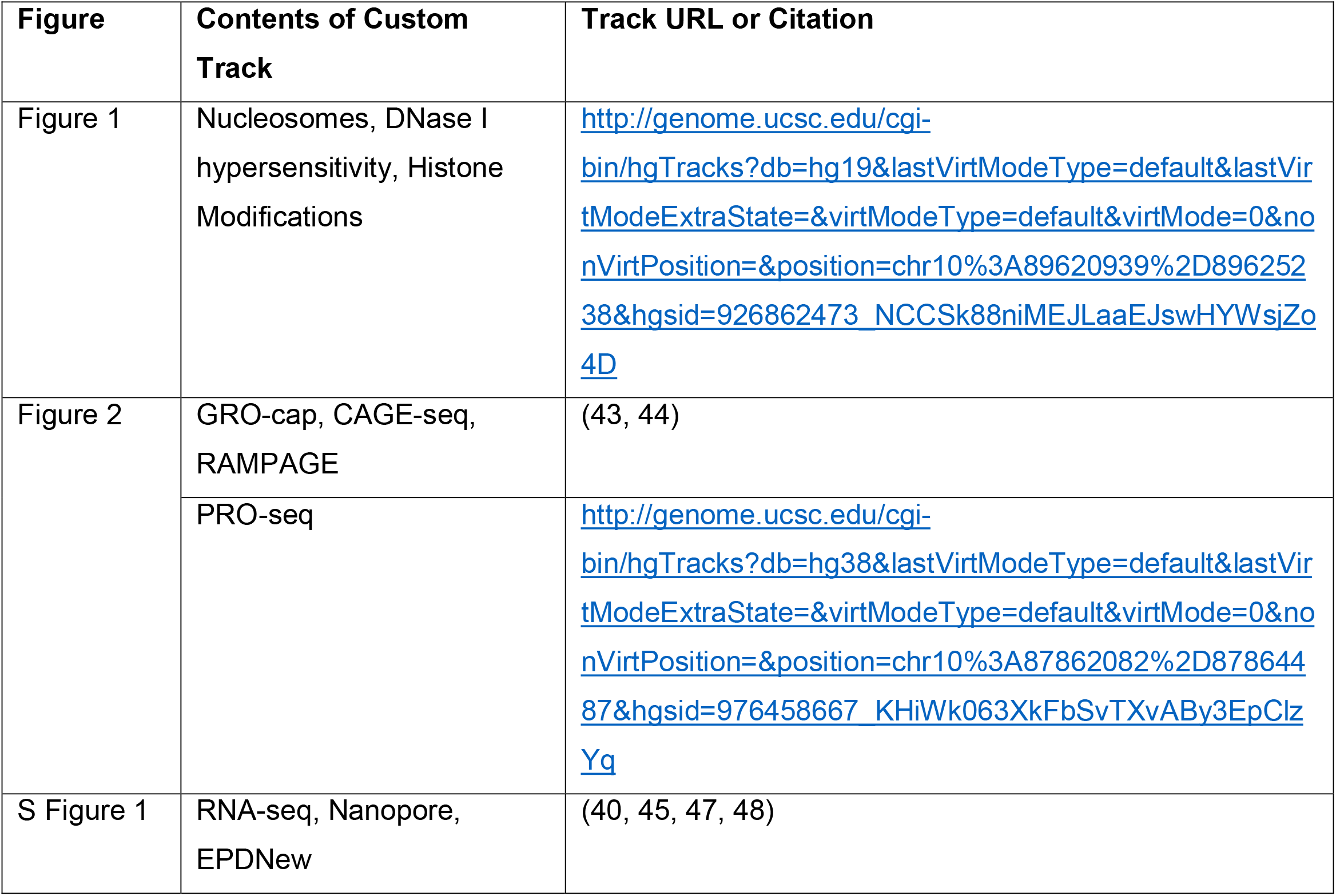

#### Gene Ontology Terms and Hierarchical Clustering Analyses

Gene ontology (GO) analysis was performed using the STRING software (https://string-db.org/), which also provided information on enriched pathways curated in Reactome (https://reactome.org), and KEGG. The GO terms associated with TFs binding the upstream region were compared to GO terms associated with TFs binding the downstream promoter region. The GO terms unique to upstream TFs were of interest to our analysis (Table 1). To contextualize the roles of the various TFs at upstream or both regulatory regions, we performed network analysis using Cytoscape 3.7.0 (https://cytoscape.org/) on the interaction data generated by STRING software (https://string-db.org/).

Ontology Terms Associated with Upstream *PTEN* Promoter Transcription Factors.

**Table 1: Ontology Terms Associated with Upstream *PTEN* Promoter Transcription Factors**
| **Ontology Term** | **Term ID** | **Description** |
| --- | --- | --- |
| **Molecular Function**<br>**GO Term** | GO:0004722 | protein serine/threonine phosphatase activity |
|  | GO:0008138 | protein tyrosine/serine/threonine phosphatase activity |
|  | GO:0033293 | monocarboxylic acid binding |
| **Biological Process**<br>**GO Term** | GO:0006953 | acute-phase response |
|  | GO:0008283 | cell population proliferation |
|  | GO:0050771 | negative regulation of axonogenesis |
|  | GO:0034121 | regulation of toll-like receptor signaling pathway |
|  | GO:0006954 | inflammatory response |
|  | GO:0007169 | transmembrane receptor protein tyrosine kinase signaling pathway |
|  | GO:0050670 | regulation of lymphocyte proliferation |
|  | GO:0030856 | regulation of epithelial cell differentiation |
|  | GO:0071230 | cellular response to amino acid stimulus |
|  | GO:0007259 | receptor signaling pathway via JAK-STAT |
|  | GO:0008406 | gonad development |
|  | GO:0008637 | apoptotic mitochondrial changes |
|  | GO:0032663 | regulation of interleukin-2 production |
|  | GO:0090329 | regulation of DNA-dependent DNA replication |
|  | GO:0035265 | organ growth |
|  | GO:2000378 | negative regulation of reactive oxygen species metabolic process |
|  | GO:0048483 | autonomic nervous system development |
|  | GO:0090102 | cochlea development |
|  | GO:0010677 | negative regulation of cellular carbohydrate metabolic process |
|  | GO:0043388 | positive regulation of DNA binding |
|  | GO:0010470 | regulation of gastrulation |
|  | GO:1905330 | regulation of morphogenesis of an epithelium |
|  | GO:0048813 | dendrite morphogenesis |
| **Cellular Component**<br>**GO Term** | n/a | n/a |
| **KEGG Term** | hsa01521 | EGFR tyrosine kinase inhibitor resistance |
|  | hsa05323 | Rheumatoid arthritis |
| **RCTM Term** | n/a | n/a |

## RESULTS

### Public genome browser data suggest novel regulatory region for PTEN with promoter characteristics

To initially screen the *PTEN* locus (Figure 1A) for novel regulatory elements identifiable via NGS technologies, we overlaid tracks in the UCSC Genome Browser against transcriptional landmarks upstream of the *PTEN* CDS. We began by visualizing stranded poly(A)+/-RNA-sequencing results from whole cell and nuclear lysates of ENCODE Tier One and Two cell lines (Figure 1B) to comprehensively assess all transcripts initiated near the *PTEN* locus. Alignments of RNA transcripts that extended toward *PTEN* (including both poly-adenylated and non-poly-adenylated) were concordant with previous promoter sequences, identified as multiple reads with 5’ ends in the Stambolic/Sheng and Han regions. The GC-rich nature of *PTEN*’s 5’UTR likely affected the read depth of the region between the Stambolic and Han regions, as indicated by the lack of RNA transcript depth in the most GC-dense sequences and no splice donors or acceptors exist in the region (Figure 1B). Interestingly, RNA-seq reads were also identified approximately 800bp upstream of the full-length promoter (Figure 1B). The presence of sequence reads 5’ of the canonical promoter on the plus strand suggests the possibility of upstream transcription initiation in the direction of *PTEN*.

We further validated the pattern we analyzed using the Eukaryotic Promoter Database (EPDnew) and long-read RNA-sequencing by Nanopore. The EPDnew algorithm bioinformatically predicts promoter sequences using high-throughput promoter mapping experiments and NGS data (47, 48). This algorithm predicted a promoter sequence about 1.1kb upstream of the putative *PTEN* promoter, which coincides with the ENCODE RNA-seq track data showing potential upstream transcription initiation (Supplemental Figure 1). Additionally, comparison of NanoPore long-read RNA-seq and the stranded RNA-seq tracks show similar trends in transcription initiation location and read depth (Supplemental Figure 1).

In conjunction with our assessment of transcripts via ENCODE RNA-sequencing data, we next investigated the chromatin landscape near *PTEN* to analyze the regulatory potential of the upstream region. We first looked for evidence of open chromatin conformations, suggesting DNA accessibility to transcription factors (TFs) by nucleosome depletion and DNase hypersensitivity (DHS) sequencing data available in ENCODE. Nucleosome packing data was available from micrococcal nuclease sequencing (MNase-seq) on ENCODE Tier 1 cell lines GM12878 and K562 cells (41). These data demonstrate that not only are nucleosomes depleted in the Sheng and Han regions, but are also depleted near the E-box domain and in the upstream region of our interest highlighted above (Figure 1C). Similar patterns of DNA accessibility are observed in DNAse hypersensitivity sequencing (DNase-seq) experiments sourced from the Roadmap Epigenomics Consortium (49). DNase-seq results from numerous cell lines and tissue types indicate that DNA in the Sheng region and our upstream region of interest represent euchromatin-dominant regions (Figure 1C). Due to the importance of open chromatin for transcription factor (TF) binding and transcription initation, these data support the presence of RNA transcripts near and upstream of the canonical *PTEN* promoter.

Since we observed transcripts upstream of the canonical *PTEN* promoter and the chromatin landscape in the region suggested it may be accessible to TFs, we next analyzed histone marks that may predispose the region to promoter characteristics. Additionally, we sought to understand how our findings may be influenced by the E-box domain located at −2180 to −2175, which may have enhancer-like effects on *PTEN* transcription (34). To answer these questions, we aligned the promoter-associated histone modifications H3K27Ac and H3K4Me3 and the enhancer-associated histone modification H3K4Me1. H3K27Ac and H3K4Me3 modifications have been observed in numerous tissue types surrounding the upstream and Sheng regions, a histone pattern generally observed in active promoter regions (40). Our analyses showed H3K4Me1 modifications flanking the H3K4Me3 marks around the upstream regulatory region in a nested manner, further supporting the existence of an upstream promoter (50). Therefore, both the Sheng region and upstream regulatory region maintain stronger promoter characteristics than enhancer characteristics (Figure 1C).

### PTEN Upstream Regulatory Region contains functional TSRs

Our initial screening of the *PTEN* locus using publicly available NGS data (Figure 1) showed us that a novel region upstream of the canonical promoter generated transcripts in the direction of *PTEN*, was depleted of nucleosomes, and maintained promoter-associated histone modifications. These findings pushed us to search for exact transcription start sites within this upstream promoter. Therefore, we scanned the established and the potentially novel *PTEN* promoter regions for evidence of TSSs using various NGS tools designed to determine the 5’ end of mRNA transcripts. Global run-on sequencing of transcripts with 5’ 7-methyl guanosine residues (GRO-cap) and cap analysis gene expression sequencing (CAGE-seq) datasets were initially examined to compare where transcripts are initiated at the *PTEN* locus (Figure 2A). GRO-cap is most useful when paired with CAGE-seq to capture both nascent, potentially fleeting transcripts, in addition to stable transcripts that are likely to be translated (43). Upon analysis of GRO-cap data from GM12878 and K562 cells, we observed that nascent *PTEN* transcription (elucidated by peaks in the track related to signal intensity) occurs not only in the Sheng minimal promoter, but also focuses on two sequences of the upstream region we identified in RNA-seq (Figure 2A). Negligible transcription initiation is observed at the homologous sequence of the Han constitutive mouse promoter. Additionally, the CAGE-seq tracks show that capped transcripts can be captured and sequenced to the two upstream promoters and the canonical promoter regions (Figure 2A). These tracks supported our original RNA-seq findings that both the Sheng and upstream regions generate RNA capable of translocation from the nucleus.

**Figure 2:**
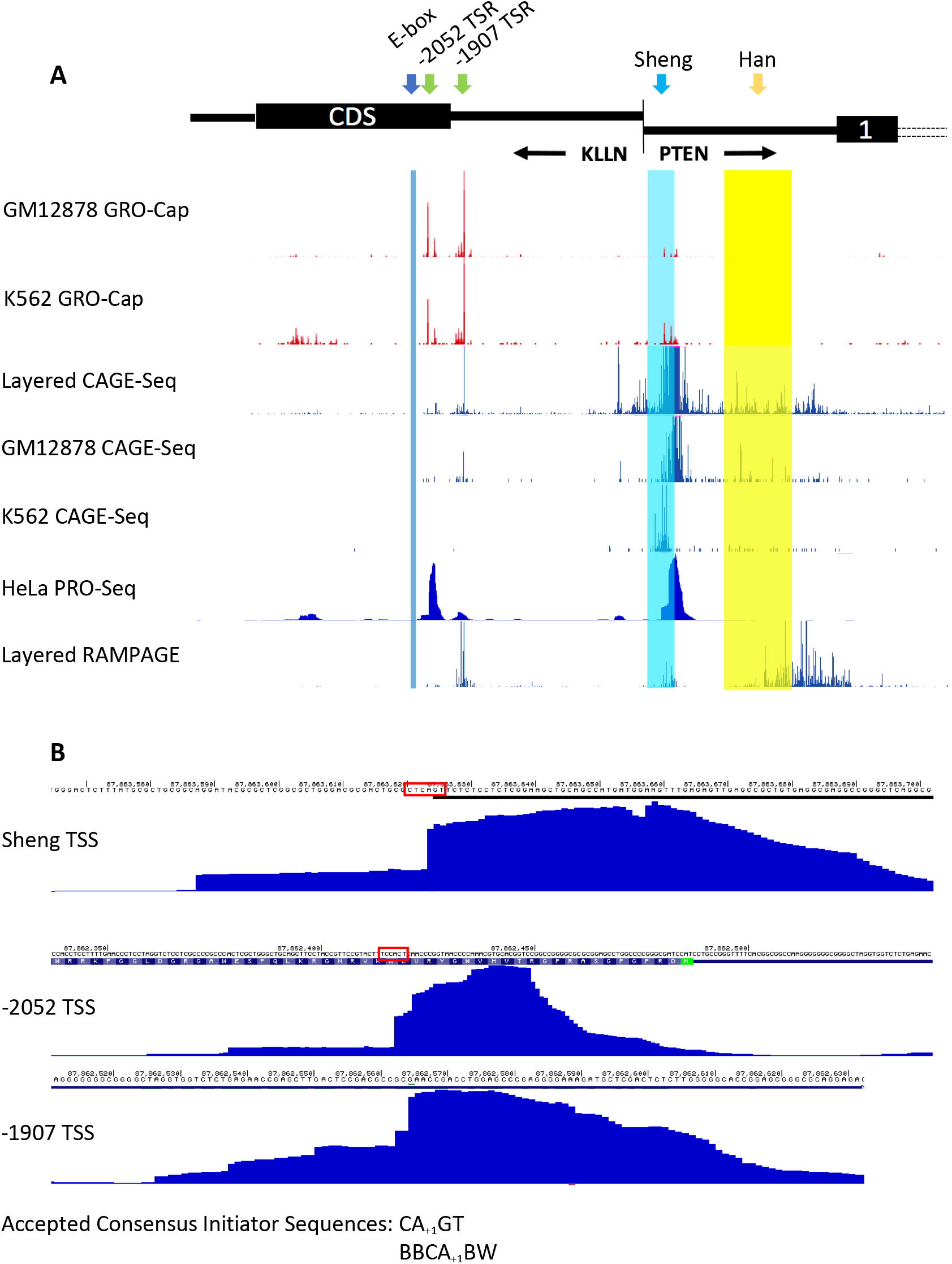
Novel transcription start regions identified upstream of canonical *PTEN* promoter. (A) Multiple Run On Sequencing tracks corroborate transcription initation at canonical promoter and upstream locations at *PTEN* locus. Global run on sequencing of 5’ 7-methyl guanosine residues (GRO-cap) tracks sourced from Human GRO-cap Hub in GM1878 and K562 (43). Cap analysis of gene expression sequencing (CAGE-seq) tracks visualized as layered track of 7 ENCODE cell lines and from GM12878 and K562 cells (44). Precision run on sequencing (PRO-seq) track from HeLa cells visualizes amplified regions of Pol II binding and thus transcription initiation sites (23). RNA annotation and mapping of promoters for the analysis of gene expression (RAMPAGE) sourced from 7 ENCODE cell lines (44). (B) Initiation site from PRO-seq analyses (23) suggest start site at 87,863,624 for canonical *PTEN* TSS, 87,862,418 for furthest-upstream novel TSS, and 87,862,565 for second novel TSS. The canonical TSS mirrors the consensus initiator sequence of BBCA_+1_BW, supporting the PRO-seq data. The −2052 TSS differs from the consensus initiator by one base pair. The −1907 TSS does not contain a canonical Inr sequence but utilizes a noncanonical Inr to begin transcription.

**Figure 3:**
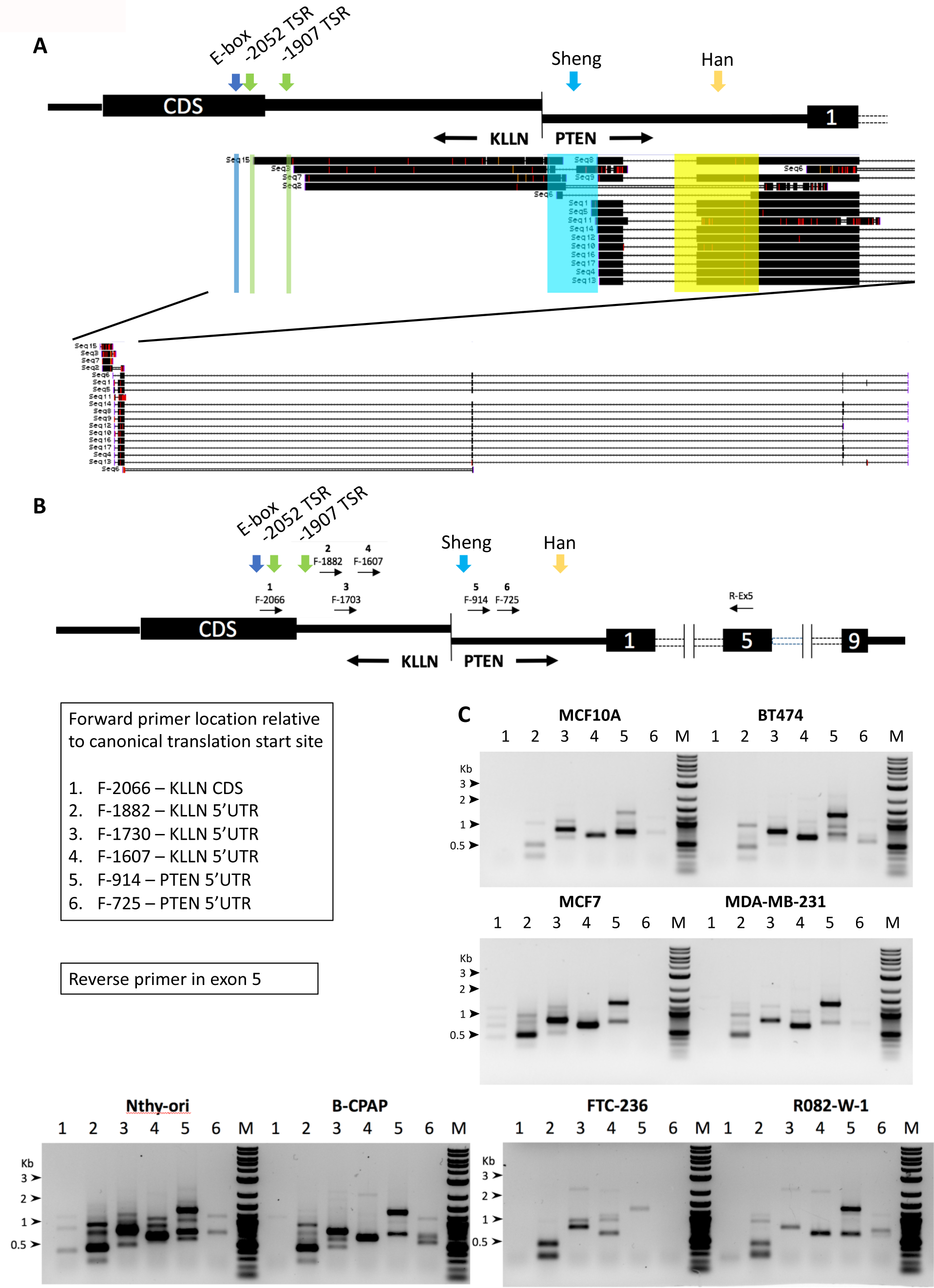
Extended *PTEN* transcripts identified in lymphoblast, thyroid, and breast cell lines through reverse transcription assays. (A) *PTEN* transcripts isolated from 5’Rapid Amplification of cDNA Ends Sequencing (5’RACE-Seq) in lymphoblast cell line (LCL) clones. Visualized are 17 sequences collected from clones, with the 5’ reverse primer located within exon 4 of *PTEN*. Four of the 17 sequences amplified through the canonical promoter and into the upstream region. (B) Schematic of RT-PCR primers chosen to isolate canonical and potential novel upstream *PTEN* transcripts. RNA from human breast cell lines was converted to cDNA These forward primer sequences were paired with a reverse primer at Exon 5 of *PTEN*. (C) *PTEN* transcripts identified in four breast cell lines and four thyroid cell lines using aforementioned primer pairs. Similar patterns of transcript amplification observed throughout all eight cell lines, with slight variability in transcript abundance based on cell line. Multiple bands observed in many of the lanes, suggesting that alternative splicing of these *PTEN* transcripts is possible.

Since initial Run On Sequencing data support the existence of two upstream promoters for *PTEN*, we sought to map the transcription initation to a specific initiator (Inr) sequence. To do so, we visualized Precision Run On Sequencing (PRO-seq) and RNA annotation and mapping of promoters for analysis of gene expression (RAMPAGE) from HeLa cells and a layering of seven cell lines, respectively (23, 44). We were therefore able to directly compare the DNA sequence of canonical and upstream initation sites to the consensus Inr sequences of CA_+1_GT (23) and BBCA_+1_BW (51), where B represents an A, C, or T and W is an A or T (Supplemental Figure 2). An exact Inr sequence was found to match the PRO-seq peak within the Sheng promoter of CTCA_+1_GT, with the +1 site 846 bp upstream of the canonical ATG translation start codon (Figure 2B). These data suggest that most productive transcription within the minimal promoter occurs downstream of the putative MaxTSS defined within −951 and −925. The most upstream PRO-seq peak contained the sequence TCCA_+1_CT, with the +1 location of the Inr site 2052 bp upstream of the canonical ATG within the CDS of KLLN. It is worth noting here that the upstream Inr differs in one nucleotide compared to the consensus Inr sequences. Similarly, the start site at −1907 bp upstream of the translation start site identified by our PRO-seq and other Run On track analysis, does not reveal an Inr sequence. However, the data showing initiation of transcription in these regions strongly suggest promoter charactersitics inspite of the absence of canonical Inr sequences,, which may impact the effciency of transcription initiation from this site (51). Collectively, these data support the presence of two active upstream TSRs from which extended *PTEN* transcripts are generated, albeit with lower levels of transcription than the canonical promoter. We next sought to experimentally validate and characterize *PTEN*-specific transcripts from this upstream region.

### Extended *PTEN* transcripts are generated from the suggested upstream TSRs

Based on our multifaceted and compelling NGS results, we hypothesized that if *PTEN* truly has an active upstream promoter sequence, we should be able to isolate 5’ extended *PTEN* transcripts from cell lines and human tissue samples. Therefore, we completed two preliminary analyses to isolate extended *PTEN* transcripts *in vitro*. First, we identified the 5’ ends of transcripts using 5’ Rapid Amplification of cDNA Ends (5’RACE) PCR in lymphoblast cell lines (LCLs; Figure 3A). We extracted, cloned, plated, and sequenced *PTEN* transcripts from the LCLs after 5’RACE and a secondary nested PCR and were able to identify a number of unique transcripts from the clones, which were subsequently visualized as a custom UCSC track (Figure 3A). We identified 17 unique *PTEN* transcripts among which 4 of the 17 transcripts included sequences upstream of the canonical promoter that terminate near our identified TSRs. Therefore, our data suggest that the newly identified upstream TSRs may not be the preferred *PTEN* TSRs, but nonetheless are still functional and productive (Figure 3A).

Next, we designed staggered forward RT-PCR primers upstream of the canonical promoter for testing on breast and thyroid cell lines expressing *PTEN* (Figure 3B). By amplifying *PTEN* transcripts using staggered forward primers, we were able to capture the general region in which upstream transcription is initated (Figure 3C, D). RT-PCR produced only weak amplification of transcript using the −2066F/-E5R primer pair in all four cell lines (Figure 3C, D). The forward primer utilized was 21 nucleotides, so slight amplification from the −2052 TSR may account for the weak amplification as slight overlap between the primer and TSS would allow for some transcript identification. Interestingly, we observed stronger amplification when using the −1882F, −1730F, and −1607F in all four cell lines (Figure 3C, D). These primers would only amplify if transcripts were initiated upstream of the primer sequence. Therefore, these data indicate that the TSS must lie within −2066 and −1882, which directly supports the novel −2052 and −1907 TSRs identified through NGS. Additionally, the replicable pattern of multiple bands per primer set and each cell type suggests that there may be alternative splicing of these extended *PTEN* transcripts. The −914F/-E5R primer set was designed to represent canonical transcripts, as the forward primer was set slightly downstream of the Stambolic TSR; this primer pair amplified as expected (Figure 3C, D). Notably, the −725F/-E5R primer set only weakly amplified in all four cell lines. This is likely due to inefficient reverse transcriptase enzyme function in the ∼85% GC-rich region (Figure 3C, D); low RNA-seq read depth from the NGS data corroborates this result (Figure 1B).

After isolating *PTEN* transcripts that initiated upstream of the canonical promoter in breast, thyroid, and lymphoblast cell lines, we replicated these results in primary human tissue. Using the staggered upstream forward RT-PCR primers described above (Figure 3B), we sequenced transcripts from human breast and thyroid tumor/matched normal tissue (Figure 4A). These forward primers were paired with a reverse primer at Exon 9 of *PTEN*. As observed for the breast cancer lines, no significant transcript amplification was observed with primers located at position −2066F. However, amplification was observed in all tumor and normal pairs from forward primer pairs located −1882 and downstream. Notably, multiple bands of *PTEN* cDNA were observed throughout the multiple primer pair experiments in both tumor and normal samples, suggesting that these *PTEN* transcripts were alternatively spliced. To further understand the transcripts derived from the upstream TSS, bands identified from the −1882F/-E9R primer set were extracted, cloned, plated, and Sanger sequenced (Figure 4B). Sanger sequencing confirmed the sequences to be those of *PTEN* and not the result of random mispriming (Figure 4B). Sequencing revealed that transcription initiation occurred before −1882 and could be followed by splicing at −1826 to multiple locations in the 5’UTR of *PTEN* for multiple alternative transcripts (Figure 4B). Although these splice variants do not directly affect the *PTEN* coding sequence, splicing from −1882 to −17 would eliminate uORF1 and uORF2 from the *PTEN* 5’UTR, which could impact translation of the standard PTEN protein (27). Splice donor and acceptor scores were calculated for the 5’UTR splice variants, with all three locations obtaining scores supporting the validity of splicing (Figure 4C). Additionally, two of the five types of alternative *PTEN* transcripts sequenced directly spliced out exons (Figure 4B), which, if translated, would deleteriously impact *PTEN* structure and function by deleting the PIP_2_-binding motif (52, 53). Collectively, these data show that upstream *PTEN* transcription initiation occurs in human breast and thyroid tissue, both in tumor and healthy samples.

**Figure 4:**
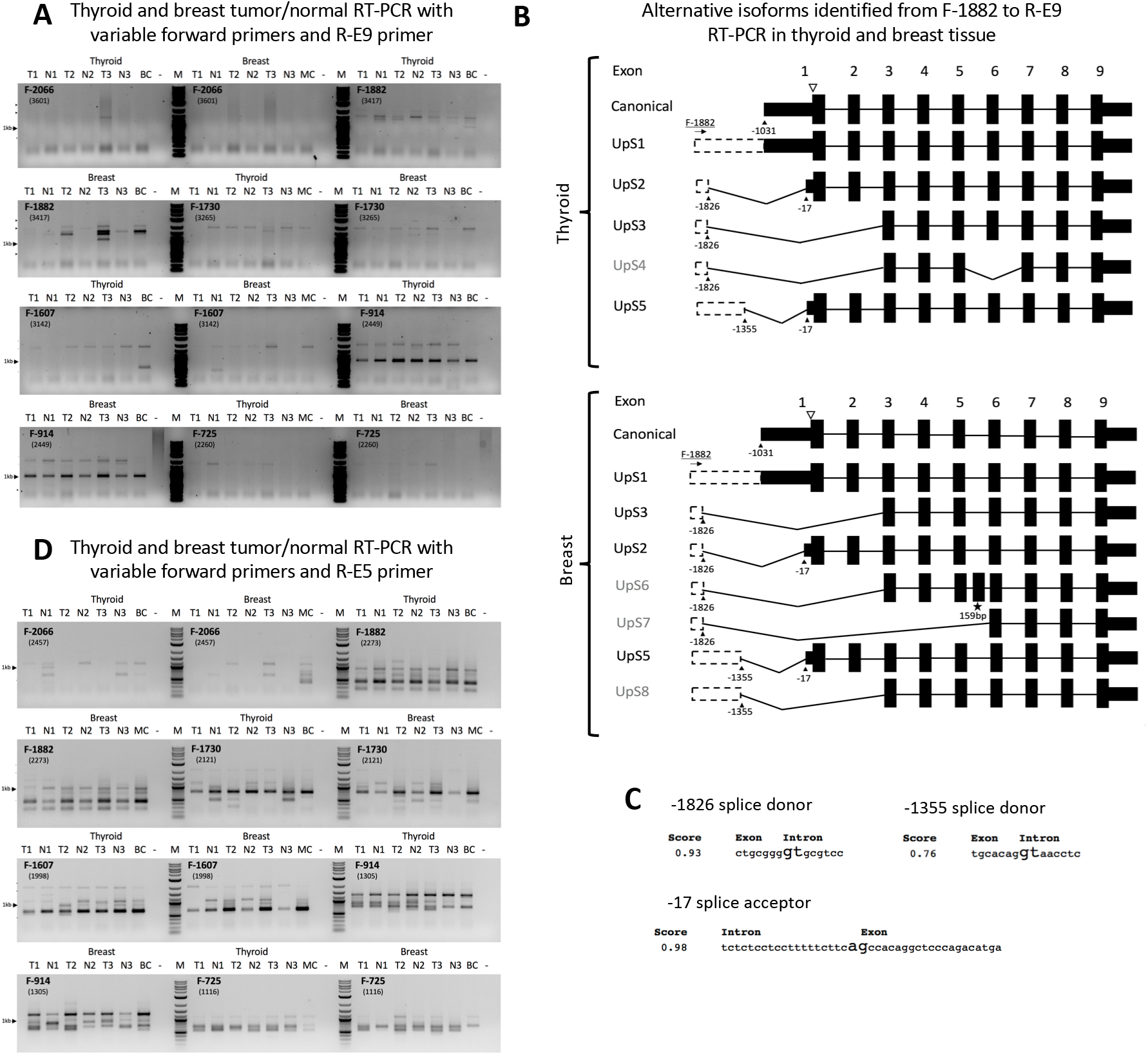
Extended *PTEN* transcripts can undergo alternative splicing in human breast and thyroid tissue. (A) *PTEN* RT-PCR from thyroid and breast tissues using variable forward primers defined in Figure 3B and an exon 9 reverse primer. Multiple bands per lane represent *PTEN* transcripts of different length identified within the same sample. Darker banding observed for shorter reverse primer samples than longer samples, suggesting that the shorter reverse primer increased yield. Positive control lanes use B-CPAP cells to represent thyroid and MCF7 cells to represent breast cancer. Negative controls lanes run with vehicle. Note that some primer sets show relatively even band intensities across all samples while others (i.e. *PTEN*_-1882F) show a marked difference between tumor and normal, especially for breast. (B) Most common *PTEN* splice variants isolated from breast and thyroid cancer and normal tissue RT-PCR with −1882F/-E9R after bacterial transformation. Frequency of clones containing splice variant recorded. UpS2 is the most common splice variant in Thyroid-derived clones, while UpS2 is the most common variant in breast-derived clones. (C) Splice donor fidelity scores identified for three most prevalent donor sites calculated by NNSPLICE v0.9 (Berkeley Drosophila Genome Project). Higher fidelity scores predict successful splicing events with 1.00 representing a perfect score. Fidelity scores presented represent strong splicing scores, supporting the sequences’ potential to act as a splice donor/acceptor. (D) *PTEN* RT-PCR from thyroid and breast tissues using variable forward primers defined in Figure 3B and an exon 5 reverse primer. Experimental setup mimics prior data shown in Figure 4A except the reverse primer; reverse primer changed to capture wider variety of splice variants.

### TFs unique to the upstream promoter may drive tissue- or cell-specific expression of extended *PTEN transcripts*

After validating the concordance between our observations utilizing the public NGS data and our in-house experiments supporting the presence of two novel upstream promoter sequences for *PTEN*, we compiled TF-binding data from ENCODE and from available literature. We first compared TF binding between the upstream promoters versus the canonical promoters using chromatin immunoprecipitation sequence data (ChIP-Seq) on 340 TFs from 129 cell types from ENCODE, which is the most comprehensive single source of TF binding available (45) (Figure 5A). We found that most TFs were shared by both regions (Supplemental Table 1; Figure 4B). Interestingly, we found a larger number of TFs that were unique to the upstream promoter regions (38 TFs) compared to those binding uniquely to the canonical Sheng minimal promoter (3 TFs) (Supplemental Table 1).

**Figure 5:**
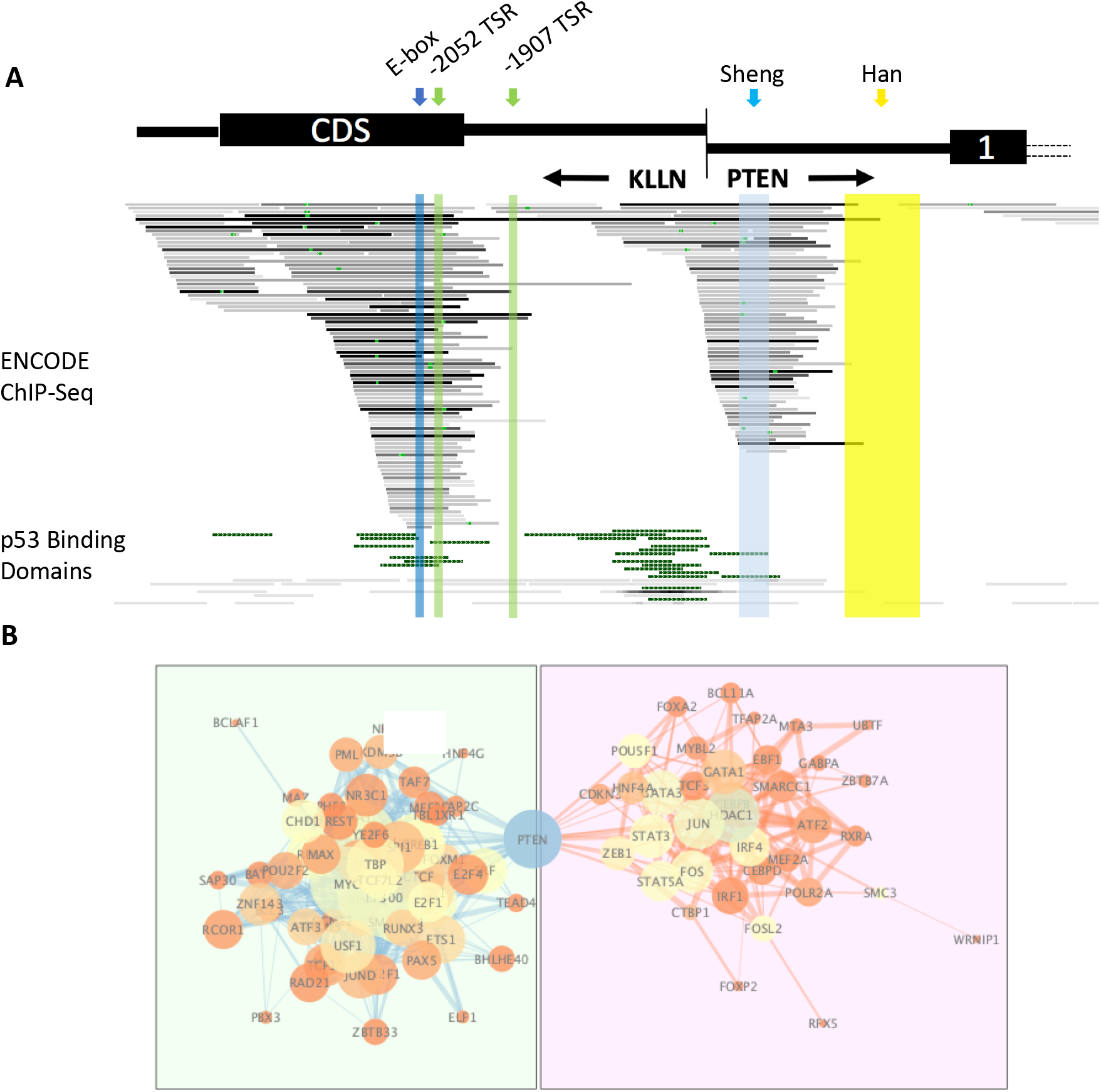
Upstream *PTEN* promoters regulated by unique transcription factors compared to canonical promoter. (A) ENCODE 3 ChIP-Seq data sourced from UCSC Genome Browser representing TFs binding at *PTEN* locus from 91 cell lines (45). Binding region and affinity for each individual TF marked by single horizontal line, with increase in darkness representing stronger affinity. P53 binding domains from ENCODE also presented (45). (B) Hierarchical clustering of Gene Ontology Pathway Analysis connections for TFs that uniquely interact with the upstream *PTEN* TSRs. Clusters integrated through *PTEN* interaction, with TFs binding to the canonical promoter initiation complex (PIC) or the upstream PIC shown on left and TFs binding uniquely to the upstream PIC to the right. Coolness of the node represents betweenness centrality, and thickness of the edge represents repeatability and prevalence of the finding.

To ascertain the overarching biological pathways associated with the TFs specific to the *PTEN* locus, we utilized a Gene Ontology enrichment analysis to identify the GO terms associated specifically with upstream-binding TFs. These associated GO Terms, including Reactome and KEGG annotations as well, are included in Table 1. Interestingly, enriched GO terms for the upstream promoters suggested involvement of inflammatory processes. Upstream-promoter TFs associate with enriched terms such as acute-phase response, inflammatory response, regulation of toll-like receptor signaling pathway, and regulation of lymphocyte proliferation among anothers, which may indicate that the upstream promoters drive function-specific or cell-specific expression of *PTEN* isoforms (Table 1).

After completing the GO, KEGG, and Reactome term analyses, we assessed the interactions of the upstream TFs via pathway analysis (Figure 5B). By using PTEN as a central node to mediate interactions between the TFs, we visualized the relationships between unique upstream TFs and TFs shared between the upstream and canonical promoters. Cooler colored nodes reveal increased betweenness centrality, suggesting that these nodes mediate a larger proportion of the interactions in our analysis. Notably, HDAC1, JUN, and FOS were predicted to bind uniquely to the upstream TSRs and appear to mediate regulation of these TSSs, while TBP, MYC, USF1, and other TFs previously associated with *PTEN* interact at all TSRs (Figure 5B). Collectively, our study shows that additional promoter regions exist −2052 and −1907 upsteam of the canonical and currently recognized PTEN promoter. Further, these upstream TSRs have a unique TF profile compared to the canonical promoter, potentially supporting *PTEN’s* inflammatory role.

## DISCUSSION

This study represents a comprehensive reappraisal of the architecture of the upstream regulatory region of *PTEN*, which was accomplished by leveraging public NGS data from ENCODE. Inspection and analysis of chromatin structure, histone modifications, and stranded RNA-sequencing from ENCODE datasets (40–42) confirmed prior work on the minimal promoter (36) and suggested that another upstream location harbors a transcription regulatory region (Figure 1). These initial data indicated the possibility of an additional upstream *PTEN* promoter or a broader TSR that may drive expression of diverse *PTEN* transcripts. Subsequently, through a combination of publicly available NGS data and in-house experiments, we demonstrated that there are two novel *PTEN* TSRs upstream of the Sheng promoter (Figure 2,3,4). These novel *PTEN* TSRs, located within the KLLN CDS and 5’UTR, are identifiable by an Inr sequence at −2052 and a non-canonical Inr at −1907, respectively (Figure 2) (23, 51, 54, 55). Moreover, the upstream TSRs maintains a unique TF profile compared to the Sheng promoter (Figure 5). Due to the differences in TF binding between the regions, unique expression patterns may be driven from the upstream TSRs. This finding corroborates the long-standing claim that genes often have multiple TSRs that can be regulated in tissue- and/or function-specific manners (22, 23, 56).

Interestingly, the Sheng promoter represents the only *PTEN* TSR residing within the currently accepted full length promoter: −1344 to −745 (32). Although previous studies have hinted at upstream regulation of *PTEN* (34), −1344 is the furthest upstream sequence currently assessed in patients with PHTS or PHTS-like phenotypes. Because of this definition, if mutations outside this region impact *PTEN* transcription regulation, then the latter are not analyzed in a CLIA/CAP setting. Considering that between 15-75% of CS/CS-like patients and 40% BRRS patients patients do not have germline *PTEN* mutations (12), other perturbations to *PTEN* expression must play into the development of these phenotypes. It has previously been established that even slight decreases in TSG gene dosage can drive pathologic effects (18, 19, 57). Therefore, we stress that a stronger understanding of *PTEN* transcriptional regulation may help to explain the molecular underpinnings of PHTS-like phenotypes. Beyond our promoter’s potential impact on understanding PHTS-like phenotypes in the future, identification of the −2052 TSR remaps the 5’UTR of *PTEN*. The edge of the 5’UTR is no longer directly opposing the *KLLN* 5’UTR, which may impact how the regulation of *KLLN* is assessed, as the identifeid *PTEN* 5’UTR from this report now overlaps the *KLLN* CDS (Figure 1A). We included the murine constitutive *Pten* promoter (38) in our reappraisal, but our NGS data suggest that it does not maintain function in humans. Throughout our multiple analyses, no specific TSR could be visualized (Figures 1, 2A). Therefore, the Han murine *Pten* promoter acts as a reference point in our study with other known *PTEN* promoters.

Although our study is heavily predicated on findings catalogued in public databases, it offers a basis for future research studying the molecular and clinical importance of these novel upstream *PTEN* TSRs. It was beyond the scope of this study to fully characterize the molecular characteristics and activity of the novel *PTEN* TSRs, but we have presented validation experiments that isolate extended transcripts from multiple cell lines and human tissue samples. The −2052 Inr extends the 5’UTR for *PTEN*, introducing multiple strong splice donors and thus increasing the potential for alternative splicing. By increasing the splicing opportunities for *PTEN* transcripts through the addition of strong splice donors and acceptors, more complex regulation can occur. Strong splice donors exist at −2041 and −1826, with the most common splice acceptor located at −17 (Figure 4). Interestingly, a splice acceptor also exists before exon three of canonical *PTEN*, which is the next acceptor utilized if the −17 acceptor were skipped. Use of this splice donor would remove the first and second exon, thus eliminating the PIP_2_-binding motif and part of the dual-specificity phosphatase domain (52, 53). The loss of this domain would functionally limit the lipid phosphatase activity of any PTEN protein translated from these transcripts. Additionally, changing the 5’UTR of a transcript can greatly impact the feasibility of its translation, as adding nucleotides increases the likelihood of secondary structure impeding translation (58). This is especially significant considering the GC-richness of the *PTEN* 5’UTR. Further, alternative splicing of extended *PTEN* transcripts from − 1826 to −17 would remove upstream open reading frames (uORFs) known to generate unique isoforms of *PTEN* (27). With these findings in mind, future research may look to identify how activation of the upstream promoters results in changes to the *PTEN* transcript population available for translation. The implications of these findings in cancer are still not clear as transcript splicing was found to be similar in both cancer and normal tissues tested (Figure 4). However, expanding study to other PTEN-associated overgrowth syndromes or ASD models may elucidate specific splicing trends from upstream transcripts.

In addition to identifying new active *PTEN* TSRs, we also present a comprehensive analysis of TF binding at the *PTEN* locus (Figure 5, S Table 1). We identified region-specific TF binding for the *PTEN* locus, which may offer alternative processes for tissue specific or disease state specific regulation of *PTEN* expression (Figure 5). Due to the lower resolution of ChIP-seq, we were unable to compare TFs unique to each of the upstream TSRs; however, estimating differences between the upstream and canonical regions was feasible. By leveraging Gene Ontology enrichment analysis, we were able to connect the TF profile of upstream transcription initiation with certain molecular function and biological processes, (Figure 5B, Table 1). Interestingly, immune-related biological processes were enriched for the grouping of TFs that bind only upstream, suggesting that *PTEN* may undergo specific transcription activation from upstream sites under conditions of inflammation. PTEN is intricately involved in both central and peripheral immune regulation mechanisms which may underly *PTEN*-associated overgrowth syndromes (59–63) and explain its association with ASD (64, 65). Involvement of the *PTEN* upstream promoter in immune dysregulation may be interesting and warrants further investigation.

Complicating the TF binding profile and alternative splicing patterns of the *PTEN* promoter is identifying how each of the upstream TSRs drive expression patterns given their unique characteristics. GRO-cap, CAGE-seq, and RAMPAGE show stronger signals near the −1907 PRO-seq peak than at the Inr at −2052 (Figure 2). These datasets show that transcription initiation can occur from either of the TSRs, but due to their conflicting strength between experiment types, we cannot determine if one TSR is consistently dominant over another or how each is utilized. Specifically, GRO-cap and CAGE-seq data suggest that the −1907 TSR generates more nascent transcription and capped transcripts than the −2052 TSR. However, the −2052 TSR with a canonical initiator sequence is stronger in PRO-seq. Differences in cell type and experimental design may explain these findings. Interestingly, the −1907 TSR does not contain a canonical Inr (Figure 2B) but still maintains active transcription. Around 45% of Inr’s do not match the BBCA_+1_BW consensus sequence (51), so it is not beyond reason that *PTEN* transcription initiates here based on the observed behavior of the RNA polymerase. Ultimately, our 5’RACE sequencing and RT-PCR experiments (Figure 3,4) also support that transcripts can initiate at both locations, which externally validate that both upstream TSRs are active.

In addition to the findings presented, we wish to bring attention to this work as a roadmap for future research utilizing publicly available genomics data. If a commonly studied TSG like *PTEN* has a previously undefined regulatory region, it is possible that unknown TSRs for other clinically relevant genes may be clarified through the leverage of publically-available NGS data. Mechanizing sinks of public Run On or gene expression data offers a low barrier to entry for researchers attempting to develop hypotheses without the capability to complete these technically complicated and cost-challenging experiments themselves. However, these data should not be the only source of information used to categorize a region as transcriptionally active; hypothesis-driven experiments that validate *de novo* NGS findings are crucial. Because of our initial findings regarding these *PTEN* promoters, wider translational questions can now be asked. For instance, mutation-negative individuals with PHTS-like phenotypes may be subject to sequencing of a redefined full *PTEN* promoter to identify links between our novel TSRs and pathology.

Altogether, this study demonstrates the utility of publicly-available NGS data in refining our regulatory understanding of a particular gene. These data are consistent with our validating in-house experiments and have helped redefine the architecture of the *PTEN* locus. Our identification of two upstream TSRs indicates a possible need to expand clinical sequencing approaches for individuals who are otherwise *PTEN* mutation-negative. Ultimately, this effort may be an important step toward illuminating the murky ambiguous genetic origins of disease in PHTS-like *PTEN* mutation-negative individuals.

## AVAILABILITY

All genome browser track data have been linked above (Methods UCSC Tracks Table).

## ACKNOWLEDGEMENT

We would like to thank Tammy Sadler for helpful discussions. CE is the Sondra J. and Stephen R. Hardis Endowed Chair of Cancer Genomic Medicine at the Cleveland Clinic, and is an ACS Clinical Research Professor.

## FUNDING

This work and open access fee was supported, in part, by the Sondra J. and Stephen R. Hardis Endowed Chair in Cancer Genomic Medicine (to C.E.).

## CONFLICT OF INTEREST

All authors declare no relevant conflicts of interest.

## Notes

### Competing Interest Statement

The authors have declared no competing interest.

