## Supplemental Materials for "Redefining the *PTEN* Promoter: Identification of Two Upstream Transcription Start Regions"

**S1**



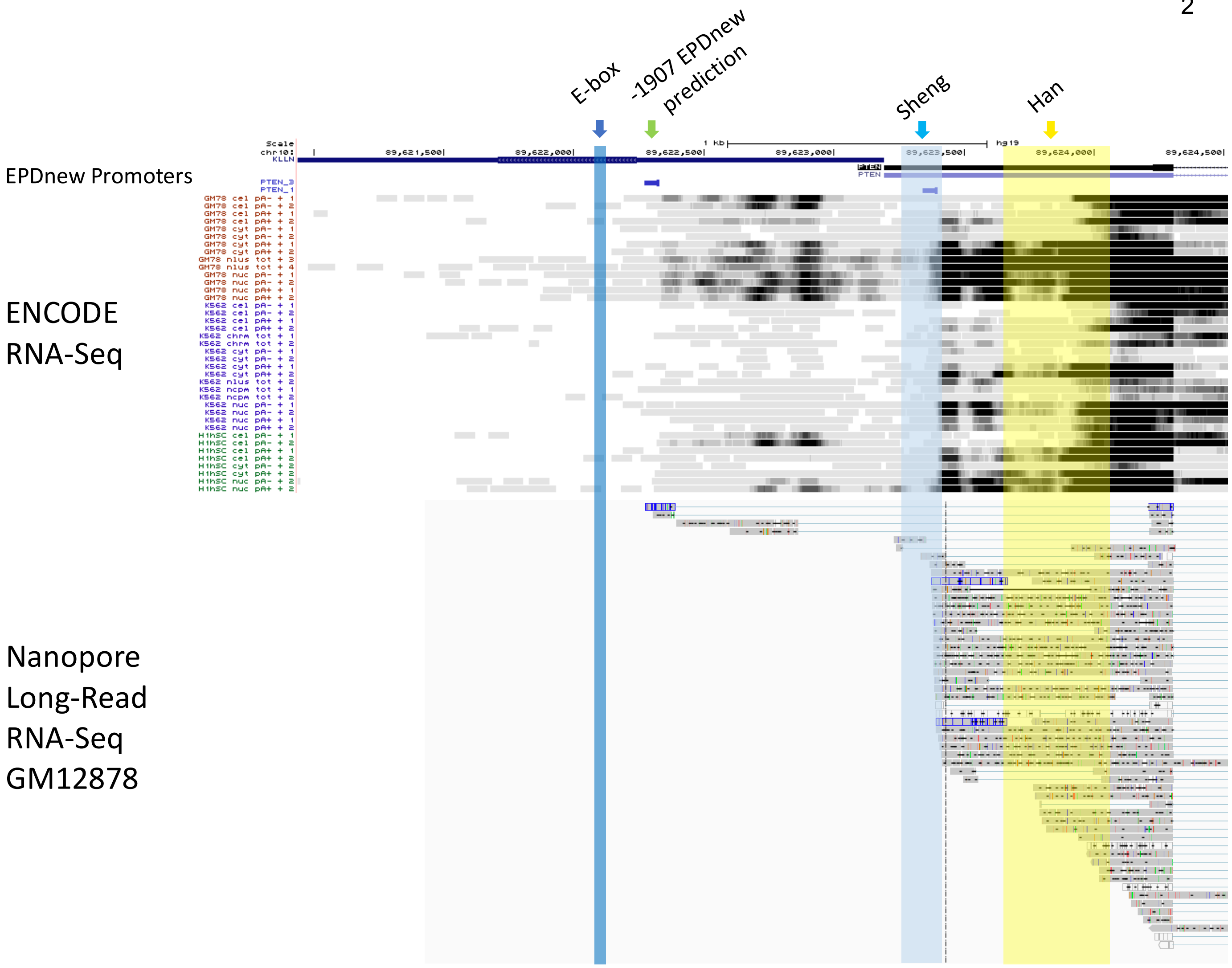


**Supplementary Figure 1: EPDnew promoter prediction and Nanopore long-read RNA-sequencing data suggest upstream *PTEN* TSR.** EPDnew promoter prediction overlaid with ENCODE 3 RNA sequencing and Nanopore long-read RNA sequencing. The EPDnew promoter prediction algorithm predicts a promoter sequence at 1907 and 838 bases upstream of the PTEN translation start site. Nuclear, nucleolar, and cytoplasmic poly(A)+/- RNA sequencing coincides, as transcripts are identified initiated in all cell lines studied at this location and upstream of it. Further, Nanopore long-read sequencing of GM12878 cells also shows transcription of *PTEN* initiating upstream of the Sheng promoter.

S2



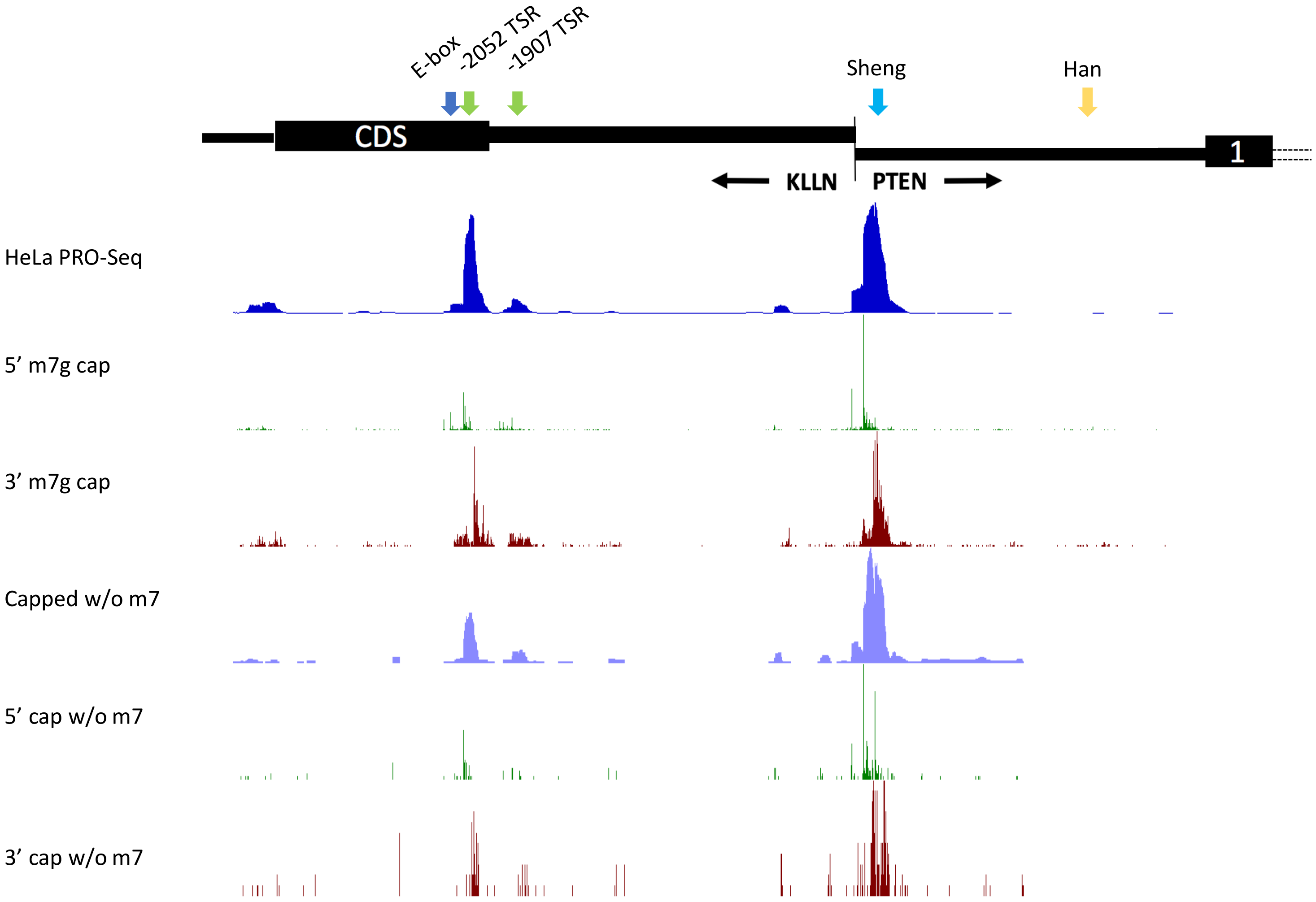


**Supplementary Figure 2: PRO-seq tracks support extended *PTEN* transcripts initiated upstream of Sheng promoter.** Precision Run-On sequencing (PRO-seq) on HeLa cells (23) identifies two TSRs upstream of canonical promoter. PRO-seq tracks represent TSRs, with designation made between cap methylated and non-methylated transcripts (w/ m7g versus w/o m7g). 5’ and 3’ tracks refer to the Transcription Start Site (TSS) and Transcription Pause Site (TPS), respectively, for both cap methylated and nonmethylated transcripts. Strong PRO-seq signal for methylated transcripts at 2052, 1907, and 846 bp upstream of the canonical PTEN translation start indicate three active TSRs for the *PTEN* locus.

S. Table 1: **Transcription factors binding at PTEN locus by cell type and binding location**

| **Transcription Factors** | **Cell Types (from ENCODE ChIP-Seq)** | **Binding location (Upstream = U, Canonical = C)** |
|---|---|---|
| ATF2 | GM12891 | U |
| ATF3 | GM12892 | U/C |
| BATF | H1-hESC | U/C |
| BCLAF1 | | U/C |
| BCL11A | H54 | U |
| BCL3 | HCT116 | U/C |
| BHLHE40 | HEK293T | U/C |
| CCNT2 | | U/C |
| CEBPB | OCI-LY1 | U |
| CEBPD | | U |
| CHD1 | OCI-LY7 | U/C |
| CHD2 | | U/C |
| CREB1 | Parathyroid adenoma | U/C |
| CTBP1 | Raji | U |
| CTCF | SH-SY5Y | U/C |
| E2F1 | WERI-Rb-1 | U/C |
| E2F4 | WI38 | U/C |
| E2F6 | adrenal gland | U/C |
| EBF1 | astrocyte of the spinal cord | U |
| EGR1 | body of pancreas | U/C |
| ELF1 | breast epithelium | U/C |
| ELK1 | cardiac muscle cell | U/C |
| EP300 | choroid plexus epithelial cell | U/C |
| ETS1 | epithelial cell of proximal tubule | U/C |
| FOS | fibroblast of pulmonary artery | U |
| FOSL2 | fibroblast of villous mesenchyme | U |
| FOXA1 | foreskin fibroblast | U/C |
| FOXA2 | foreskin keratinocyte | U |
| FOXM1 | gastroesophageal sphincter | U/C |
| FOXP2 | | U |
| GABPA | keratinocyte | U |
| GATA1 | liver | U |
| GATA3 | mammary epithelial cell | U |
| GRp20 | | U |
| GTF2F1 | | U/C |
| HDAC1 | omental fat pad | U |
| HDAC2 | ovary | U/C |
| HMGN3 | | U/C |

| | | |
|---|---|---|
| HNF4A | spleen | U |
| HNF4G | stomach | U/C |
| IRF1 | uterus | U |
| IRF4 | | U |
| JUN | | U |
| JUND | | U/C |
| KAP1 | | U |
| KDM5B | | U/C |
| MAX | | U/C |
| MAZ | | U/C |
| MEF2A | | U |
| MEF2C | | U/C |
| MTA3 | | U |
| MXI1 | | U/C |
| MYBL2 | | U |
| MYC | | U/C |
| NR3C1 | | U/C |
| NRF1 | | U/C |
| PAX5 | | U/C |
| PBX3 | | U/C |
| PHF8 | | U/C |
| PML | | U/C |
| POLR2A | | U |
| POU5F1 | | U |
| POU2F2 | | U/C |
| RAD21 | | U/C |
| RCOR1 | | U/C |
| RELA | | U/C |
| REST | | U/C |
| RFX5 | | U |
| RUNX3 | | U/C |
| RXRA | | U |
| SAP30 | | U/C |
| SIN3AK20 | | U/C |
| SIN3A | | U/C |
| SMARCB1 | | U/C |
| SMARCC1 | | U |
| SMC3 | | U |
| SP1 | | U/C |
| SP4 | | U |
| SPI1 | | U/C |

| | | |
|---|---|---|
| SRF | | U/C |
| STAT3 | | U |
| STAT5A | | U |
| TAF1 | | U/C |
| TAF7 | | U/C |
| TBL1XR1 | | U/C |
| TBP | | U/C |
| TCF3 | | U |
| TCF12 | | U/C |
| TCF7L2 | | U/C |
| TEAD4 | | U/C |
| TFAP2A | | U |
| TFAP2C | | U/C |
| TFAP4 | | U |
| UBTF | | U |
| USF1 | | U/C |
| USF2 | | U/C |
| WRNIP1 | | U |
| YY1 | | U/C |
| ZBTB33 | | U/C |
| ZBTB7A | | U |
| ZEB1 | | U |
| ZNF143 | | U/C |
| p53 | | U/C |
| EGR-1 | | U/C |
| CBF1 | | Not Assigned |
| PPARg | | Not Assigned |
| NFKB | | Not Assigned |
| SNAIL | | Not Assigned |
| EVI1 | | Not Assigned |
| NOTCH1 | | Not Assigned |
| SALL4 | | Not Assigned |
